# AlphaFold2 enables accurate deorphanization of ligands to single-pass receptors

**DOI:** 10.1101/2023.03.16.531341

**Authors:** Niels Banhos Danneskiold-Samsøe, Deniz Kavi, Kevin M. Jude, Silas Boye Nissen, Lianna W. Wat, Laetitia Coassolo, Meng Zhao, Galia Asae Santana-Oikawa, Beatrice Blythe Broido, K. Christopher Garcia, Katrin J. Svensson

## Abstract

Secreted proteins play crucial roles in paracrine and endocrine signaling; however, identifying novel ligand-receptor interactions remains challenging. Here, we benchmarked AlphaFold as a screening approach to identify extracellular ligand-binding pairs using a structural library of single-pass transmembrane receptors. Key to the approach is the optimization of AlphaFold input and output for screening ligands against receptors to predict the most probable ligand-receptor interactions. Importantly, the predictions were performed on ligand-receptor pairs not used for AlphaFold training. We demonstrate high discriminatory power and a success rate of close to 90 % for known ligand-receptor pairs and 50 % for a diverse set of experimentally validated interactions. These results demonstrate proof-of-concept of a rapid and accurate screening platform to predict high-confidence cell-surface receptors for a diverse set of ligands by structural binding prediction, with potentially wide applicability for the understanding of cell-cell communication.

## Introduction

Many secreted proteins, polypeptides, and peptides constitute signaling molecules that control intercellular communication by binding and activating membrane receptors^1,2^. Upon receptor binding, these molecules directly coordinate short or long-distance signaling responses and biological functions such as cell growth, survival, and metabolism^1,3,4^. The human secretome contains at least 2,000 secreted proteins, not counting posttranslationally processed fragments and peptides^5^. The vast majority of these ligands have no assigned function or cognate receptor. Single-pass transmembrane receptors, also known as bitopic proteins, represent more than 50 % of all transmembrane proteins^6^ (1300 proteins in humans^7^) and include receptor tyrosine kinases (RTKs), cytokine receptors, enzymes, and extracellular matrix proteins^4,8–10^. Surprisingly, most ligands for these receptors remain unknown. Deorphanization of functional receptors can open up entirely new fields in biology and offer new therapeutic avenues^11^.

Performing experimental screens to identify ligand-receptor pairs is challenging for several reasons. Mapping interactions at the cell surface is inherently more difficult than identifying intracellular interactions. This is because extracellular ligand-receptor interactions often have low affinity and fast dissociation rates, making high-throughput screening methods such as affinity purification challenging^12,13^. Similarly, binding screens using an individual ligand applied to a receptor in solution are time-consuming, not applicable for all receptor types, and may lack the cellular environment necessary for posttranslational modifications or co-receptor binding^13,14^. Lastly, cell-based CRISPR screens are limited by the ability to gain sufficient receptor expression and the lack of expression of essential coreceptors^13,15^.

With the revolutionizing ability to predict protein 3D structures from their amino acid sequences, AlphaFold (AF2) has become an omnipresent tool in the field of structural biology^16,17^. AF2 can predict protein-protein interactions including heterodimeric protein complexes^18–20^, but accurately modeling membrane-spanning protein complexes can pose a challenge requiring knowledge of topology^21,22^. Prediction of protein-protein interactions (PPIs) using AF2 have been previously benchmarked^23^. These studies have revealed that complexes originating solely from eukaryotes are predicted more accurately compared to mixed complexes, that modeling success rate varies depending on the annotated function of chains^23^, and that homomers more accurately model interfaces compared to heteromers^24^. Importantly, previous benchmark studies have not compared the use of full canonical protein sequences compared to cleaved and binding parts of structures^18,22,23,25,26^. They have also not addressed the success rate of predicting dimers that are part of larger protein complexes. This is important for screening purposes as computational cost as well as feasibility is aggravated by modeling complex size, In addition, single-pass receptors typically form homo or heteromers and ligands frequently dimerize. Receptor and ligand binding partners forming a binding complex are typically not known in the discovery phase. Previous benchmarking studies have also not addressed how false positive rate relates to ranking. In a screening setting, false positives are of less importance compared to ranking. Illustrating the discrepancy between model success and ranking, AF2 has been evaluated for screening purposes of multi-pass receptor-peptide interactions finding that protein interaction metrics can be effectively used to rank predictions. Interestingly, though success rate, as defined by DockQ score, was acceptable or high for all eleven tested peptides, ranking varied from 1-25^22^ preventing its use as a classifier.

Here, we benchmark AF2 as a screening tool for single-pass receptors. We show that accuracy in a screening setting is dependent on complex type and cannot be inferred from general benchmarking of AF2. Specifically, we find that ranking accuracy is likely superior for ligand-single-pass receptors compared to ligand-G-protein coupled receptor (GPCR) interactions. We describe the computational and input settings for the prediction screen, performance, success rate, show that “promiscuous” ligands with many putative false predictions are likely to be more successful in predicting the correct receptor with a slight loss in accuracy, and provide proof-of-principle evidence of identification of high-confidence binders. This work provides a useful resource for future investigations and is likely to be relevant to a wide variety of fields including cancer research, immunology and endocrinology.

## Results

### Improving binding prediction accuracy with domain selection

There is currently no report of AF2’s applicability as a screening tool to predict binding of an extracellular ligand to its cognate single-pass cell surface receptor. Therefore, we first aimed to test and optimize the input sequences to test the impact on interface accuracy of ligand-receptor binding predictions. Single-pass transmembrane receptors may produce spurious predictions with intertwined transmembrane, intracellular, or extracellular domains which might interfere with ligand binding prediction^21^. Consequently, this might result in inaccurate predictions for ligand binding sites. We therefore tested the effect of removing the intracellular part of the receptor on AF2 structure prediction. Prediction of ligand-receptor binding associations was performed using either the full-length receptor consisting of the extracellular domain (ECD), the transmembrane domain (TMD), and the intracellular domain (ICD), or the ECD alone. For many secreted proteins the cleavage pattern is unknown. As this has not been tested back-to-back in previous benchmarking studies, for the ligand input, we therefore tested using either the full-length ligand (secreted protein with the pro-region) without the N-terminal signal peptide, or the processed ligand cleaved from a precursor protein. Importantly, to avoid any learning-based bias by AF2, we selected ligand-receptor pairs for genes where crystal structures from the Protein Data Bank (PDB) had not been released at the point of AF2 training. For qualitative assessment of the ligand-receptor binding prediction, we used the interface template modeling (ipTM) score for modeling protein complexes where a value closer to 1 reflects a likely protein complex with a high probability of correct interface modeling, while values lower than 0.2 indicate two randomly chosen proteins^27,28^. Importantly, the ipTM score is not influenced by the size of the protein.

To examine the impact of the ligand input sequence, we designed a set of four test ligands. These ligands all possessed annotated chains according to UniProt and their respective ligand-receptor structures were not available during AF2 training (**Table S1**). The pairs were chosen by finding ligand-receptor pairs in published databases^29,30^ that did not have a structure in PDB at the point of AF2 training date cutoff (2018-04-30). The test pairs included the following: BMP10 with its receptors BMPR1A, BMPR1B and ACVRL1; AMH with its receptor AMHR2^31,32^; ALKAL1 with its receptors ALK and LTK; and the secreted antigen CD160 with its receptor TNFRSF14/HVEM. As expected, the higher the ipTM value, the more closely AF2 predictions resembled the interactions in the reference crystal structures^31,33^. This correlation is illustrated in our predictions for the BMP10-ACVRL1 (**Figures 1A-1E**) and AMH-AMHR2 pairs (**Figures S1A-S1E**). The vast majority of contacts in the crystal structure of BMP10-ACVRL1 are located between ACVRL1 residue 20-80 and BMP10 residue 240-280 (**Figure 1A**). Surprisingly, in the majority of cases, predicting the structure including the secreted ligand led to more spurious contacts compared to predicting the full ligand with only the ECD of the receptor, the secreted ligand with the receptor ECD and ICD, or the full ligand with both the ECD and ICD (**Figures 1B and 1D**). This was caused by a flip in the contact site compared to the crystal structure (**Figure 1B**) illustrating that one cannot necessarily expect more accurate predictions from input of specific binding regions. Generally for the eight ligand-receptor pairs, the highest prediction strength was observed when predicting the full or secreted ligand in combination with only the ECD of the receptor, which led to average ipTM values above 0.7 for BMP10-ACVRL1 (**Figure 1F**), BMP10-BMPR1A/B (**Figures 1G-1H**), AMH-AMHR2 (**Figure 1I**), ALKAL1-ALK/LTK (**Figures 1J-1K**), and CD160-TNFRSF14 (**Figure 1L**). Including both the ECD and ICD domains of the receptor for prediction, alongside the full ligand, consistently led to a decline in prediction accuracy. This was evidenced by median ipTM values of approximately 0.3 for AMH-AMHR2 (**Figure 1I**), ∼0.6 for ALKAL1-LTK (**Figure 1K**), close to 0.2 for BMP10-BMPR1A (**Figure 1G**), and ∼0.3 for BMP10-BMPR1B (**Figure 1H**).

**Figure 1.**
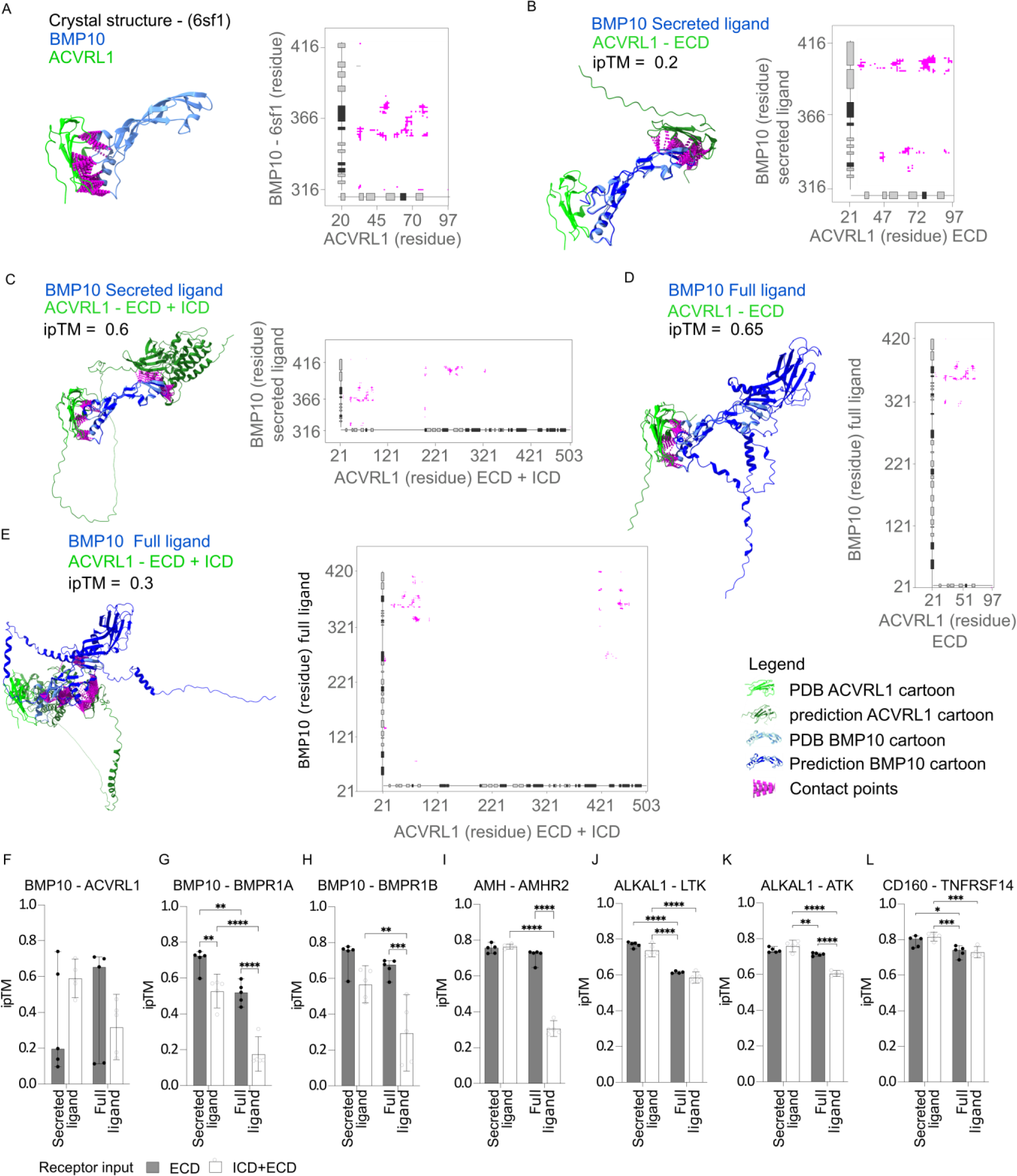
Ligand-receptor binding prediction accuracy is dependent on the sequence input. (A) PDB structure (PDB id: 6sf1) and contact map of the BMP10-ACVRL1 complex complementary to representative predictions in B-E. (B-E) Structural binding prediction and corresponding contact maps (where distances below < 8 Å were considered contacts) of ligand-receptor pairs comparing full or truncated chains of ligand and receptor. Annotation for a-e: light green cartoon: PDB database receptor, dark green cartoon: AF2 predicted receptor, light blue cartoon: PDB database ligand, dark blue cartoon: AF2 predicted ligand, magenta: contact points. Tick labels indicate residue position in full length canonical protein (BMP10 accession= O95393, AVCRL1 accession=P37023). (F-L) ipTM of the ligand-receptor pairs (F) BMP10-ACVRL1, (G) BMP10-BMPR1A, (H) BMP10-BMPR1B, (I) AMH-AMHR2, (J) ALKAL1-ALK, (K) ALKAL1-LTK, and (L) CD160-TNFRSF14 predicted by AF2 using either of the following annotated regions from UniProt: secreted ligand/chain, full ligand (pro-region and secreted ligand/chain without signal-peptide), extracellular (extracellular without signal peptide), intracellular + extracellular (full canonical sequence without signal peptide). The predictions are the median± 95% CI of five independent predictions for each ligand-receptor pair (n=5). Two-way ANOVA followed by Turkey’s test was used for multiple comparisons of differences in ipTM values between different input conditions for ligand-receptor pairs in GraphPad Prism version 9.5.0, *p < 0.05, **p < 0.01, ***p < 0.001, ****p < 0.0001.

In conclusion, selecting receptor ECDs can improve the precision of ligand to single-pass receptor binding predictions. Predicting ligand-receptor structures is not necessarily more accurate when using either the secreted or full ligand, if combined only with the receptor’s extracellular domain. On the other hand, including the intracellular domain can diminish prediction accuracy. In our limited set of test cases low ipTM values were predominately caused by spurious contact sites.

### Construction of a single-pass transmembrane receptor library

To test the performance of AF2 as a screening platform, we established a library of single-pass transmembrane proteins using sequences obtained from UniProt (**Figure 2A**). To conserve computational resources, we filtered out receptors with duplicated gene names, those without a gene name, those lacking a labeled extracellular domain, those also annotated as multi-pass, and receptors with an extracellular domain exceeding 3,000 amino acids. Since AF2 version 2.2.4 was trained on sequences longer than 15 amino acids, we also excluded entries with an extracellular domain less than 16 amino acids. This resulted in a library of 1,107 receptors. We analyzed the functional composition of the library using the membraneome database^21^, revealing that 45% of proteins are receptors, with structural/adhesion proteins at 24%, and receptor ligands/regulators at 12% (**Figure 2B, Table S2**). Single-pass transmembrane receptors span the membrane once and are classified into types I, II, II, or IV, depending on their transmembrane topology (**Figure 2C**)^7,34^. Most entries in our library are type I single-pass transmembrane receptors (86.4%), with the remaining entries being type II, III, and IV receptors, which make up 12.2%, 1.4%, and 0.1% of the library, respectively (**Figure S2A**). Receptors were distributed fairly evenly across tissues and cell types (**Figure S2B**, p < 0.001; **Figure S2C**). We explored receptor expression across tissues annotated in the Human Protein Atlas^35^, identifying that 48.5% of receptors in the library were tissue-enhanced, 25% exhibited low tissue specificity, while 0.9% were undetectable (**Figure S2D**). The library showed enrichment in cells responsive to many secreted cues, including Langerhans cells, neurons and related support cells, immune cells and enterocytes (**Figure S2E**). Further, the library was enriched for terms relating to immune function (**Figure S2F**). Taken together, the library is of use for a broad spectrum of disease areas including cancer, immunology, neurology and endocrine disorders. We also examined the ligand gene lengths for known ligand-receptor complexes^29,36^, finding that the median gene length for single-pass receptor ligands was 284 amino acids (quartiles: 189-416), while multi-pass receptor ligands were shorter at a median of 103 amino acids (quartiles: 77-152) (p<0.0001, **Figure 2D**) In summary, we show that ligand size could potentially infer receptor type, and the broad applicability of our library.

**Figure 2.**
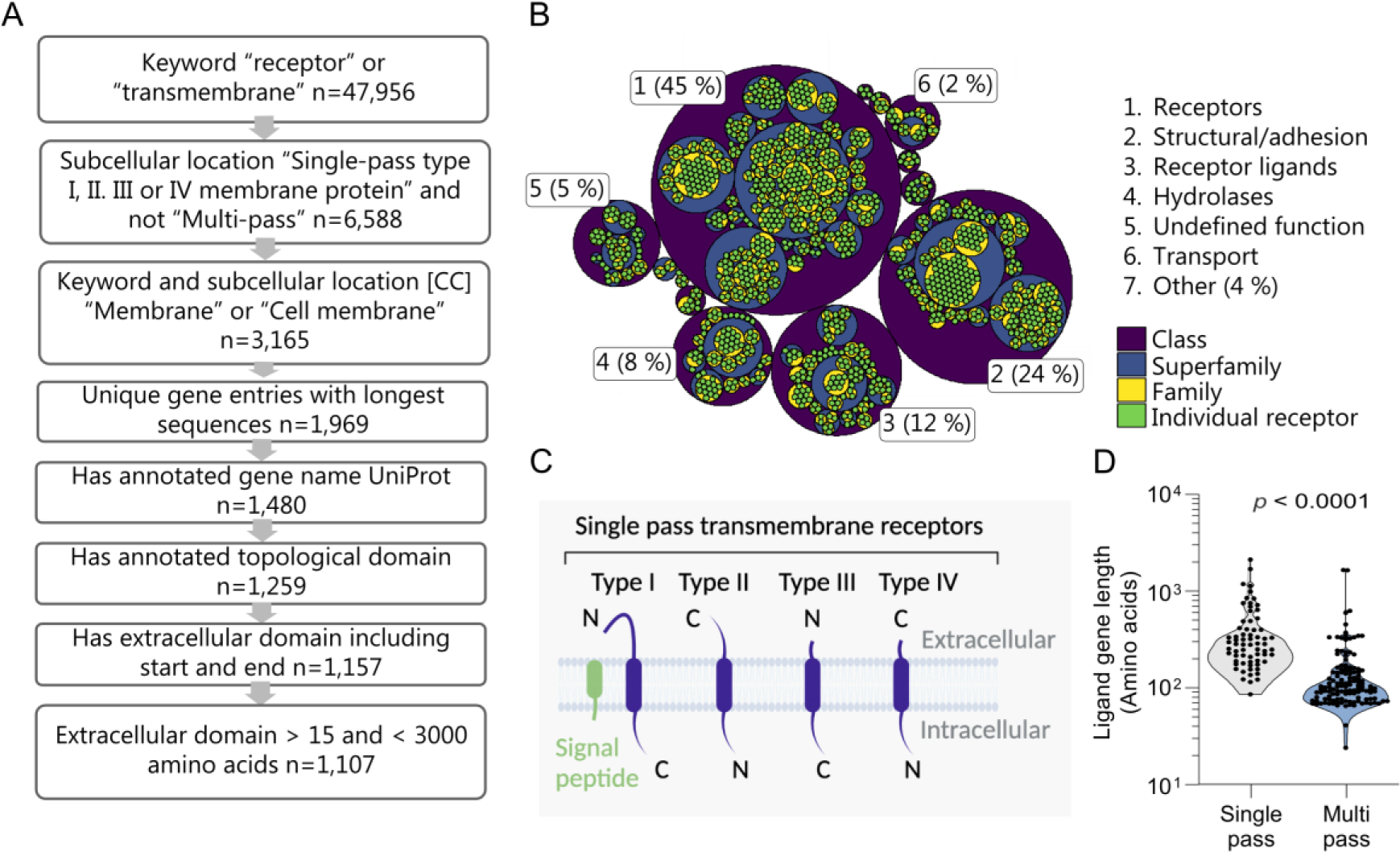
Construction and properties of a structural library of 1,107 single-pass transmembrane receptors. (A) Schematic of the receptor library construction. 1: Extract human entries with the keyword either “receptor” or “transmembrane” n=47,956. 2: Retain entries with subcellular location [CC] either “Single-pass type I, **I**, **I** or IV membrane protein” and not “Multi-pass” n=6,588. 3: Retain entries with keyword and subcellular location [CC] either “Membrane” or “Cell membrane” n=3,165. 4: Exclude duplicated gene names, for each gene retaining entries with longest sequences n=1,969. 5: Remove entries without an annotated gene name according to UniProt n=1,480. 6: Retain entries with an annotated topological domain n=1,251. 7: Remove entries without an extracellular domain including start and end n=1,157. 8: Exclude receptors with an extracellular domain shorter than 16 amino acids and longer than 3,000 amino acids, n=1,107. (B) Hierarchical classification of the receptors in the library categorized by functional annotation as defined by the membranome database as per the third of March 2023. (C) Schematic diagram of single-pass transmembrane receptor classified by type. (D) Canonical protein sequence length for ligands that bind either multi-pass or single-pass receptors expressed as amino acids (KS test p =10^−14^), n=173 multi-pass, n=64 single-pass. Significant differences in ligand length for known ligand-receptor pairs were calculated using the Kolmogorov–Smirnov test.

### Performance analysis experimentally validated ligand-receptor structures

Using the single-pass transmembrane library, we tested how accurately AF2 could predict the known receptors for ligands, establishing its effectiveness as a screening platform. We included eight receptor-ligand pairs, from previously reported datasets^29,30^, where structures were absent at the time of AF2 training for both proteins or manually curated pairs where only the receptor had an available structure^29^ (**Table S1**). Considering the often unknown processing of orphan ligands and the high ipTM values (>0.6) obtained using full ligands with the receptor’s ECD, we adopted this combination for the ligand-receptor screening (**Figure 3A**).

**Figure 3.**
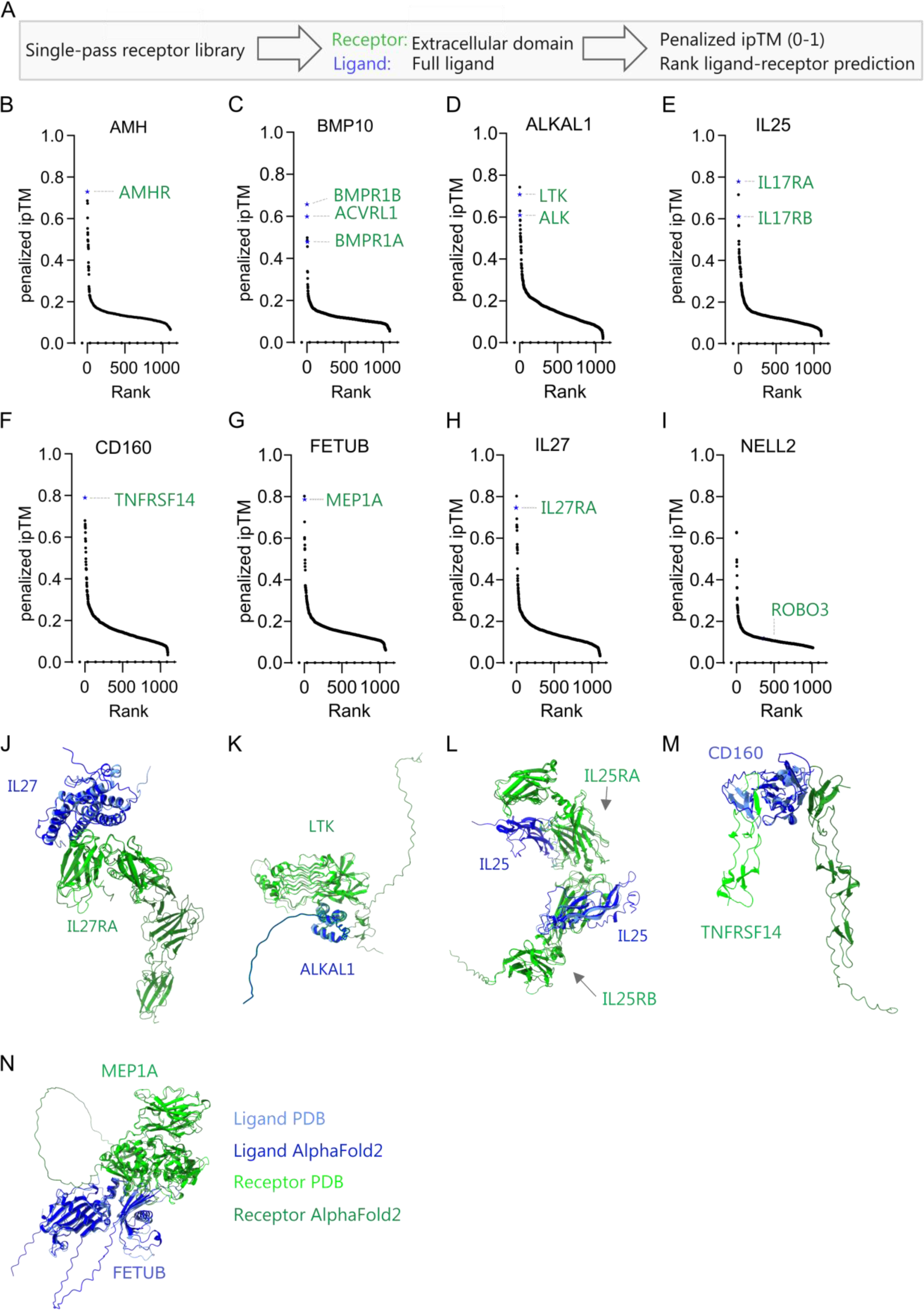
Performance and accuracy of binding prediction in a validated ligand-receptor test set. (A) Approach and metric for scoring of binding of ligands against the single-pass transmembrane receptor library. (B-I) Binding screens for test ligands. (B) Binding prediction of anti-Mullerian hormone (AMH), (C) bone morphogenic protein 10 (BMP10), (D) ALK and LTK ligand 1 (ALKAL1), (E) interleukin-25 (IL25), (F) CD160, (G) Fetuin-B (FETUB), (H) IL27, and (I) neural EGFL like 2 (NELL2) to the receptor library. Values are expressed as ranked penalized ipTM. The predictions are the median minus median absolute deviation of five independent predictions for each ligand-receptor pair. Computation was insufficient to predict structures for AMH-DCHS1 (n=1106-1107). (J-N) Representative structural binding prediction of ligand-receptor pairs comparing the PDB structures with AF2 (light blue: ligand in PDB, light green: receptor in PDB, dark blue: predicted ligand, dark green: predicted receptor) for (J) IL27-IL27Ra, (K) ALKAL1-LTK, (L) IL25-IL17RA/B, (M) CD160-TNFRSF14, and (N) FETUB-MEP1A. PDB structures used: IL17-IL27Ra (7u7n), ALKAL1-LTK (7nx0), IL25-IL17RA/B (7uwj), CD160-TNFRSF14 (7msg), FETUB-MEP1A (7uai). A two-sided Wilcoxon signed-rank test was used to compare differences in ipTM between non-binders (negative) and validated binders (positive) in R version 4.2.1. *p < 0.05, **p < 0.01, ***p < 0.001, ****p < 0.0001.

To screen and rank ligands against all the receptors in the library, we used the penalized ipTM value of five AF2 predictions^37^. With one exception, the correct receptors for the eight test cases consistently ranked among the top three receptors, yielding ipTM values between 0.6-0.8 for all ligand-receptor pairs. We accurately predicted the type II receptor AMHR for the ligand AMH as the first predicted receptor (**Figure 3B**). For the ligand BMP10, its receptor ACVRL1 is ranked number two while BMPR1B and BMPR1A ranked first and sixth, respectively (**Figure 3C**). Furthermore, for ALKAL1, the known tyrosine kinase receptor LTK^38^ was predicted as the second top ranked receptor in the screen, while the other known receptor, ALK, ranked fifth (**Figure 3D**). Importantly, other RTKs displayed low ipTMs. Given that monomeric ALKAL1 is known to form a homodimer upon ALK binding^38^, these results also suggest that binding prediction is independent of conformation changes. Furthermore, the prediction correctly identified the heterometic cytokine receptors IL17RA and IL17RB for interleukin-25 (IL25)^39^(**Figure 3E**), CD160 for TNFRSF14/CD270^40^ (**Figure 3F**), the secreted metalloproteinase fetuin-b (FETUB) for meprin A (MEP1A)^41^ (**Figure 3G**) and IL27 for IL27RA (**Figure 3H**). For one ligand, neural EGFL like 2 (NELL2), we did not predict any binding (ipTM ∼ 0.11) to the receptor roundabout homolog 3 (ROBO3)^42^ (**Figure 3I and Figure S3A**). As expected, the predicted ligand-receptor interactions demonstrated correct binding sites and locations based on the established PDB structures for IL27-IL27RA, ALKAL1-LTK, IL25-IL17RA/B, CD160-TNFRSF14, and FETUB-MEP1A complexes (**Figures 3J-3N**). The ranking of receptors was not significantly different when using the average ipTM, median ipTM, penalized ipTM, or pDockQ^37,43^, demonstrating robust prediction with low variability (**Figure S3B**). Since all correct ligand-receptor pairs had ipTM values above 0.47 and, due to thinning out in hits above this value, we hypothesized that the screen could also be performed in reverse to screen ligands for specific receptors (**Figure S4A**). We constructed a ligand library and predicted ligands for receptors that were not in the PDB database at the cut-off date for AF2 training. The ligand library was generated by including entries for genes annotated in UniProt predicted to be secreted with a sequence length between 15-2000 amino acids, excluding immunoglobulins. The ligand library comprises 1,862 unique entries (**Figure S4B**). This prediction accurately identified AMH as the ligand for AMHR as the second-ranked hit (**Figure S4C**), IL27 as the first ligand for IL27RA (**Figure S4D**), ALKAL1 and ALKAL2 as the top two ligands for LTK (**Figure S4E**), and FETUB as the top ligand for MEP1A (**Figure S4F**). In conclusion, we show that we can rapidly and reliably use AF2 as a screening method to identify ligand-receptor pairs for a diverse set of established ligand-receptor pairs with a success rate of > 85 %.

### Performance using experimentally validated extracellular protein interactions

Since our initial dataset was of limited size and since false predictions are hard to disprove, we utilized data derived from experimentally validated interactions. Since there are no available comprehensive binding datasets on secreted ligand-receptor pairs, we used a dataset of extracellular adhered protein pairs in the immunoglobulin superfamily (IgSF) identified using an ELISA-based screening platform in Wojtowicz *et al*^44^. We filtered this dataset for 83 proteins which had at least one interaction reported in both Wojtowicz et al and other studies, and thus represented high-confidence interactions. Proteins had been tested for interaction in an all-again-all manner and IgSF-IgSF pairs, here termed ligand-receptor pairs, without any documented binding thus had a low likelihood of binding. We replicated the screen using AF2, predicting all 83 proteins against each other yielding 3,401 possible combinations (**Figure 4A**). We confirmed a clear relationship between experimental binding and the AF2 predicted ipTM value (**Figure 4A**). We predicted binding, as determined by a penalized ipTM value > 0.5, in 46 % (24/52) of the experimentally validated binding pairs and 50% (42/83) of proteins ranked the correct binding partner ≤ 3 (**Figure 4B**). Similarly, we predicted binding in 60 % (12/20) of structures released after AF2 training not reported in Wojtowicz (**Figure 4A**). The screen also confirmed that pairs with a high ipTM value and a rank between one and four are likely to be binding (p < 0.01, Dunn’s test) (**Figure S5A**). A few failed binding pairs still ranked among the top three predictions for a given ligand, underscoring that prediction- and screening success are not completely correlated (**Figure 4B**). Of note, predicting reported binding pairs using AF version 2.3.1, which included PDB structures up to 2021-09-30, only rescued 3 out of 25 interactions with a penalized ipTM value < 0.5 using AF version 2.2.4 (**Figure S5B**) indicating that version 2.3.1 has a marginal effect on screen performance. Interestingly, we also noted that many proteins with an unsuccessful receptor prediction generally displayed low ipTM values (**Figures 4A and 4C**). In contrast, proteins that had at least one reported interaction, as defined by an ipTM > 0.5, in general had higher median ipTM values among top ranked non-binding receptors (**Figures 4A and 4C**). They also displayed higher median pTM, lower predicted aligned error at the interface (iPAE) and higher median ranking confidence across top ranked predictions (**Figure S5C**). Overall, we found a specificity and sensitivity similar to previously reported protein-protein predictions by AF2 (AUC = 0.769) (**Figure S5D**). However, the performance was markedly better for ligands that had over two predictions with a penalized ipTM value > 0.5 (**Figure 4D**). This came at the expense of a slightly lower accuracy in terms of rank (p < 0.05 for a correlation between number of predictions with an ipTM value > 0.5 and average rank of reported receptor with highest penalized ipTM) (**Figure 4E**). As expected, the vast majority of predictions with reported binding and a penalized ipTM value above 0.5 also accurately predicted the correct binding site and structural conformation (**Figure 4F**). Principal component analysis (PCA) of the metrics iPAE, pLDDT (ipLDDT), ranking confidence, and penalized ipTM separated binding pairs from non-binding pairs indicating that a combination of AF2 metrics can improve separation of binders from non-binders (**Figure S5E**). Closer inspection of metrics revealed that, apart from ipTM, separation of binders from non-binders were driven by iPAE (**Figure 4G**). Thus, reported binders had an iPAE significantly lower than non-binders all with a rank of ≤ 4 and ipTM>0.5 (p < 0.001) (**Figure 4G**). Setting a cut-off to the upper 95% confidence interval for iPAE of reported binders excluded 40% (25/63) of non-binding IgSFs with a rank of ≤ 4, improving specificity to 0.88 while excluding 13% (3/24) of reported binders (**Figure 4G**). In conclusion, we identify indicators of screening performance and measures to filter putative non-binders.

**Figure 4.**
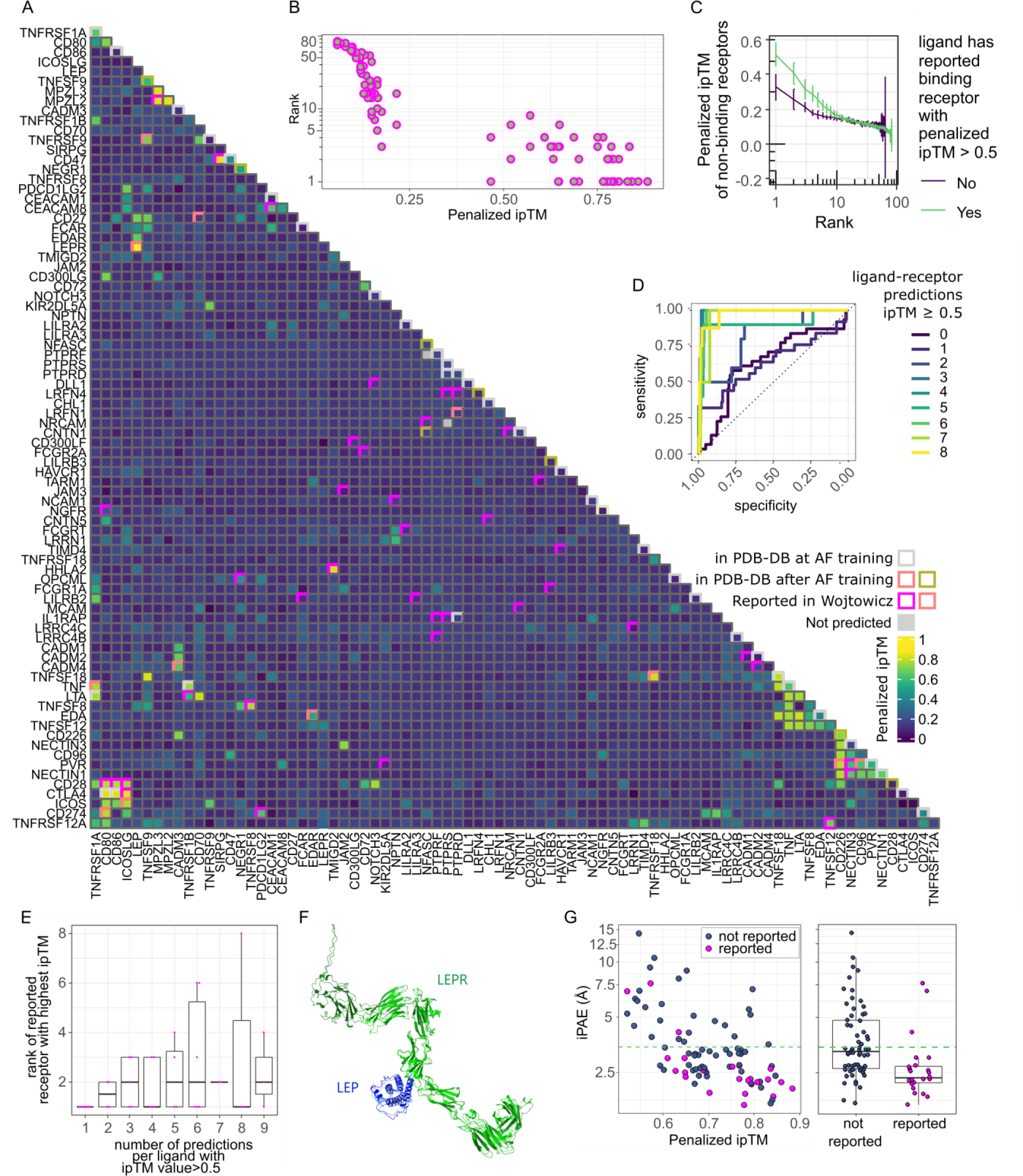
Performance using experimentally validated extracellular protein interactions. (A) Heatmap of ligand-receptor interactions that were previously experimentally tested in Wojtowicz *et al.* and elsewhere. Fill shows penalized ipTM values for ligand-receptor pairs of AlphaFold prediction. AlphaFold failed to predict any ligand-receptor pairs for the gene ‘LRFN5’ using CPUs. It was therefore dropped. Two other pairs ‘NFASC-PTPRF’ and ‘PTPRS-NRCAM’ failed prediction yielding 3,401 combinations. Gray rim: predictions that were present in the PDB database at AlphaFold training (but not identified in Wojtowicz et al), salmon pink rim: structure available in the PDB database, released after AlphaFold training date cut-off, green rim: structure released after AlphaFold training date cut-off available the PDB database, but binding not found in Wojtowicz et al., pink rim: binding reported in Wojtowicz et al. (B) Relationship between penalized ipTM value and rank for each protein for IgSF pairs reported to bind in Wojtowicz *et al*. (n=104), Spearman correlation coefficient=-0.93 (C) Penalized ipTM value as a function of rank for non-binding ligand-receptor predictions grouped by whether the ligand was successfully predicted (ipTM > 0.5) to bind its reported receptor by Wojtowicz *et al*. (n=919-2430). (D) Receiver operating curves (ROC) as a function of the number of predictions with a penalized ipTM value > 0.5. (E) rank of highest ranked binding receptor as for each ligand grouped by number of predictions where the penalized ipTM value > 0.5 (n=2-7). Spearman correlation coefficient=0.352 (p <0.05). (F) Top ranking structural binding prediction for LEP LEPR comparing the PDB structure (8avf) with AF2 (light blue: ligand in PDB, light green: receptor in PDB, dark blue: predicted ligand, dark green: predicted receptor). (G) iPAE by binding status of predictions with and ipTM >0.5 and a rank ≤ 4. Blue: not reported to bind, magenta: binding in either Wojtowicz *et al*. or pair released in PDB after AlphaFold training date cut-off (n=24-63). Boxplot displaying median, first and third quartiles and 1.5 times inter quartile range. A two-sided The Dunn’s test and the Wilcoxon signed rank test were used to compare differences in penalized ipTM and median iPAE respectively, between non-binders (negative) and validated binders (positive). P-values were adjusted for multiple testing using the Holm method in R version 4.2.1.

### Ligands that fail prediction may be inferred from AlphaFold metrics

Prediction failures are defined as known interaction partners with poor AF2 metrics (ipTM, iPAE) or incorrect structural binding sites. To understand the causes of failure, we evaluated whether the AF2 metrics could identify potential weaknesses in the predictive model. The metrics iPAE, pLDDT (ipLDDT), ranking confidence, and penalized ipTM could not distinguish reported binding-pairs that failed prediction from non-binders (**Figure S5E**). In some cases, failed predictions occurred for known interactions that rely on post-translational modifications (e.g., FCGR2A-CD72, NOTCH3-DLL1, interactions which are glycosylation-dependent). In other cases, the failed structures consisted of ligand-receptor pairs with compound interfaces (**Figure S5F**). Thus, knowledge of likely receptor formation or interface type can be useful in determining likely failures.

### Leveraging AlphaFold to identify high-confidence receptors for orphan ligands

To predict receptors for orphan secreted ligands using the single-pass transmembrane receptor library, we selected 50 ligands based on a curated library of potentially high-value orphan secreted proteins^15^ (**Table S3**). We qualitatively scored the top 20 receptors out of 1,107 receptors ranked by penalized ipTM and iPAE, taking tissue expression, known activities, and known or predicted structure into consideration (**Figure 5A**). For 90% (45/50) of ligands, we identified candidates with an ipTM value exceeding 0.5 (**Figure 5B and Table S4**), of which 18 ligands had likely receptors based on our iPAE based cut-off (**Figure 5B**). For instance, the orphan secreted glycoprotein Stanniocalcin2 (STC2) is predicted to bind to the receptor FXYD domain-containing ion transport regulator 4 (FXYD4), (penalized ipTM=0.90, iPAE=2.69) (**Figures 5B-5C**). STC2 is a regulator of calcium and is expressed broadly, whereas FXYD4 belongs to a family of proteins regulating ion transport and is exclusively expressed in the kidney^45^. STC2 binds to pregnancy-associated plasma protein-A, pappalysin-1 (PAPP-A) with structures released after AF2 training date cutoff^46^. Structurally STC2 is predicted to bind to FXYD4 in a similar position to PAPP-A (**Figure 5C**). Furthermore, we predict that the orphan ligand cerebral dopamine neurotrophic factor (CDNF), likely binds the activin receptor type-2B (ACVR2B) (penalized ipTM=0.74, iPAE=2.68) (**Figures 5B and 5D**). Structurally, CDNF is predicted to bind in the same location to ACVR2B as GDF11 (**Figure 5D**). In summary, we propose a method by which AlphaFold metrics can be used to narrowing down high-confidence receptor candidates for orphan ligands.

**Figure 5.**
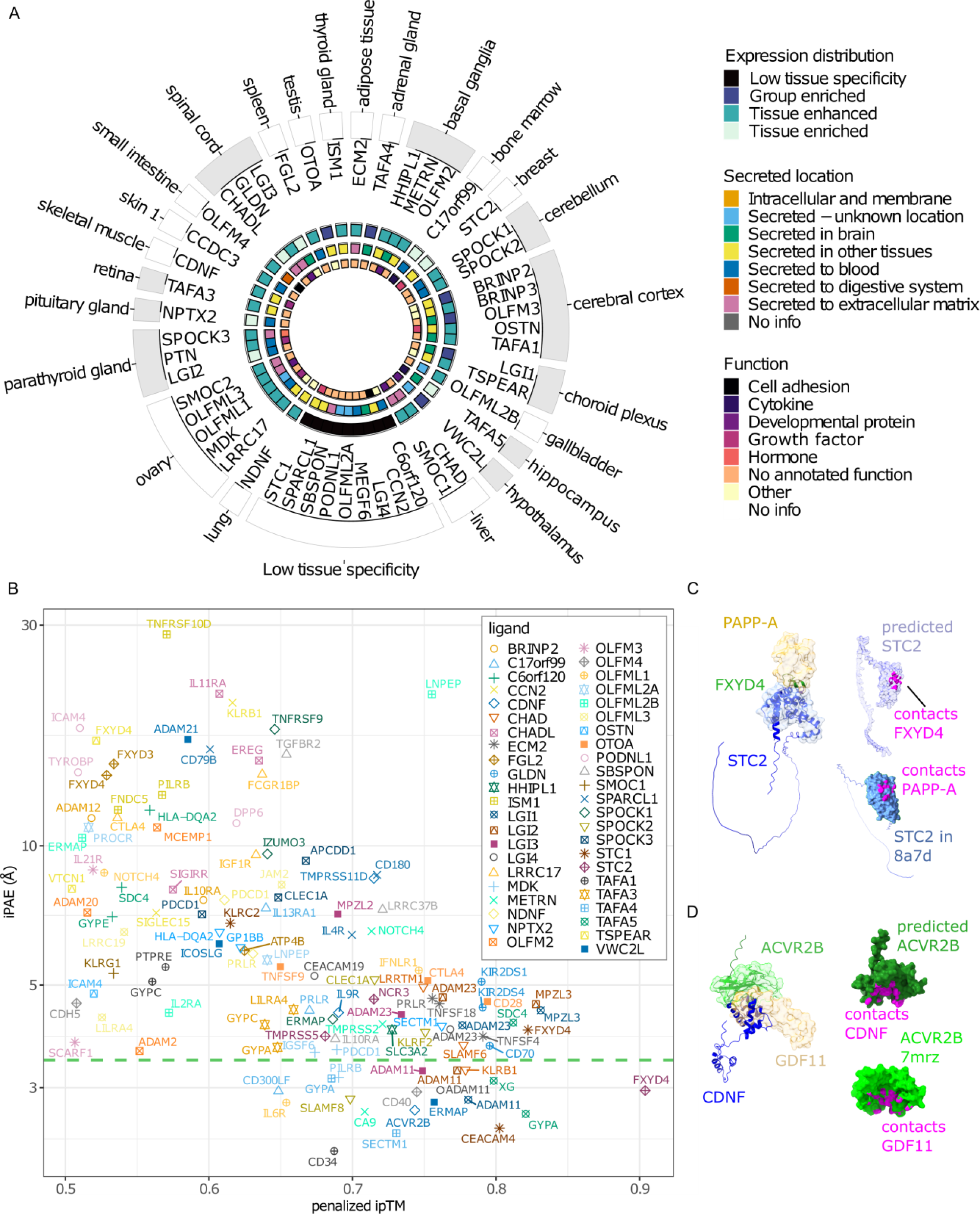
Identification of novel ligand-receptor binding pairs. (A) Schematic of orphan ligands tested using AlphaLigand. Ligands are grouped by tissue (outer text) with maximum RNA expression, inner text=gene name. Color codes going from outer to inner most ring denote: 1) gray=central nervous system, white=other. 2) Tissue distribution according to Human Protein Atlas. 3) location of secreted protein. 4) annotated function. All annotations were according to the Human Protein Atlas. (B) top one to three ranked ligand-receptor pairs with penalized ipTM > 0.5 for 45 orphan ligands (n=121). Green stippled line indicates upper 95% confidence interval of iPAE for succeeded IgSF predictions. Computational resources were insufficient to predict all five models for between 2-240 (median=19) ligand-receptor pairs across orphan ligands (Extended Data Table S4). The ipTM values of these structures were < 0.4. Based on the results from known ligand-receptor pairs, the chance of these structures ending with an ipTM value > 0.6 was below 0.1 percent. We therefore did not attempt to finish all five predictions (C) Left: representative predicted structure of the STC2-FXYD4 ligand-receptor pair matched with the partial crystal structure of STC2-PAPP-A (8a7d), blue cartoon: predicted structure of STC2, blue surface: crystal structure of STC2, green cartoon: predicted structure of the ECD of FXYD4, beige surface: crystal structure of PAPP-A. Upper right: contact points between STC2 and FXYD4, purple: predicted STC2, magenta: contact points. Lower right: contact points between STC2 and PAPPA-A, blue: STC2 (8a7d), magenta: contact points. (D) Left: representative predicted structure of the CDNF-ACVR2B ligand-receptor pair matched with the partial crystal structure of ACVR2B-GDF11 (7mrz), blue cartoon: predicted structure of CDNF, green cartoon: predicted structure of ACVR2B, green surface: crystal structure of ACVR2B, beige: crystal structure of GDF11. Upper right: Upper right: contact points between CDNF and ACVR2B, dark green: predicted ACVR2B, magenta: contact points. Lower right: contact points between ACVR2B, and GDF11, light green: ACVR2B (7mrz), magenta: contact points. Contacts were defined as van der Waals radii ≥ −0.4 Å.

## Discussion

New therapeutics are likely to target receptors or their secreted ligands^47,48^. Yet, for many hundreds of ligands identified through the secreted protein discovery initiative^48^ and in the human protein atlas secretome^5^, the receptors remain uncharacterized. In this paper, we demonstrate a simple and highly accurate screening algorithm by which AF2 can be harnessed, to predict single-pass receptors for orphan ligands. The principle that AF2 can be used to identify peptide-protein pairs has been reported^22^, but to the best of our knowledge, this is the first report documenting the use of AF2 for protein ligand-receptor interaction screening. All protein-protein interactions are not made equal. Eukaryote to eukaryote interactions in general perform better than mixed species interactions and G-protein-containing complexes perform better than other categories^23^. Here we document the performance characteristics for ligand to single-pass receptors. The finding that AlphaFold succeeds at predicting ligand-receptor binding in 46% of interactions is consistent with previous reports on eukaryote protein-protein predictions^23,49^. Based on the data presented in this paper, this resource could be expanded to include other classes of extracellular proteins, cell-surface proteins such as lipid-anchors, or pathogen proteins^36,37^. This work presents a major advance for ligand discovery where no à priori knowledge of binding sites is needed and is broadly applicable to a diverse set of secreted ligands including cytokines, hormones, receptor tyrosine kinase ligands, and proteases.

There are limitations of the method, including the computational resources needed (**Figure S6A-S6C**), and that the prediction precision and accuracy may be influenced by the binding mode and completeness of the input data. Approximately 100 single-pass receptors are missing in the library due to a lack of annotated topological domains, as well as proteins without annotated start and end domains in UniProt. These receptors could be included by inferring topological domains using computational prediction^24^. To reduce computational requirements, we limited the library to single-pass transmembrane receptors with an extracellular domain below 3000 amino acids in length. Very long sequences often fail or require extensive computation time. Moreover, glycosylphosphatidylinositol (GPI)-anchored proteins are not included because they lack a transmembrane domain. We also selected the canonical isoform of the receptors, which were not always the longest sequences. Given that the full-length ligand performed well, it is possible that the longest splice variant would perform better. Another consideration is that ligands that bind transmembrane proteins can be monomeric, dimeric, or trimeric, or require co-factors or post-translational modification for binding^50^ which is not accounted for in our binding prediction. Additionally, this approach may not be applicable to more complex receptors or ligands that interact with multiple receptors. In some cases, we were unable to predict the binding, such as for NELL2 to its receptor ROBO3. The crystal structure for NELL2-ROBO3 includes a truncated part of the ECD of the receptor which might explain the lack of binding prediction for this ligand-receptor pair^42^.

In conclusion, this research has the capacity to serve as a valuable tool for identifying previously unknown ligand-receptor pairs across a diverse range of proteins, thus opening up new possibilities for drug discovery and development.

## Acknowledgments

K.J.S. was supported by NIH grants R01DK125260, P30DK116074, American Heart Association 23IPA1042031, MCHRI, SPARK, the Weintz Family COVID-19 research fund, the Stanford School of Medicine, and the Stanford Cardiovascular Institute (CVI). K.C.G. was funded by HHMI and R01GM150125. N.B.D.S. was supported by the Carlsberg Foundation Internationalization Fellowship, Minister Erna Hamilton Grant to Science and the Arts, and the Øllingesøe Foundation. M.Z. was supported by the American Heart Association (AHA) Postdoctoral fellowship (905674). L.C. was supported by the Stanford School of Medicine Dean’s Postdoctoral Fellowship and the American Heart Association (AHA) Postdoctoral fellowship (1011077). L.W.W. was supported by the Stanford School of Medicine Dean’s Postdoctoral Fellowship. S.B.N. was supported by the Novo Nordisk Foundation (grant awards NNF20OC0059462 and NNF21CC0073729) and the Stanford Bio-X Program.

## Author contributions

Conceptualization: N.B.D.S., K.J.S.; methodology: N.B.D.S., investigation: N.B.D.S., D.K., B.B.B., G.S.O., L.C., M.Z., L.W.W., S.B.N., K.M.J., K.C.G.; supervision and funding acquisition: K.J.S.; Writing – original and revised draft: N.B.D.S., S.B.N., K.J.S.

## Competing Interests

The authors do not declare any conflicts of interest.

## Data and materials availability

All data generated or analyzed during this study are included in the manuscript, in supporting files, and at https://github.com/Svensson-Lab/AlphaLigand. Further information and requests for resources and reagents should be directed to and will be fulfilled by the Lead Contacts, Niels Banhos Danneskiold-Samsøe and Katrin J. Svensson.

## Methods

### Construction of a single-pass transmembrane receptor library

To construct a library of single-pass receptors we searched UniProt human entries for keywords with the terms “receptor” or “transmembrane” available by 11-11-2022 (n=47,956). From this pool, we retained entries stating either “Single-pass type I, **I**, **I** or IV membrane protein” and not “Multi-pass” under subcellular location [CC], n=6,588. As we restricted our library to secreted proteins, we only retained entries with keyword and subcellular location [CC] either “Membrane” or “Cell membrane”, n=3,165. Next, in the case of duplicated gene names, we prioritized reviewed entries. In case all entries with a duplicated gene name were unreviewed we prioritized the entry with the longest sequence, n=1,969. Shorter sequences were generally truncated versions of the longest FASTA sequence. Due to limited computational resources, we restricted the library to the canonical gene sequence by UniProt, avoiding other isoforms. To further limit computational requirements, we removed entries without an annotated gene name, without an annotated topological domain, and without an extracellular domain including start and end according to UniProt, retaining 1,157 receptors. Finally, as AlphaFold was trained on sequences longer than 15 amino acids we filtered receptors with extracellular domains equal to or shorter than this. To limit computation, we also excluded entries with extracellular domains longer than 3000 amino acids retaining 1,107 receptors in the final library (**Table S2).**

### Expression of receptors across human tissues

To determine the transcriptional distribution of single-pass transmembrane receptors, we extracted RNA expression of the 1,107 entries in the single-pass receptor library in all 54 tissues and all 79 single cell types found in the Human Protein Atlas (version 23.0)^51^. Here we introduced a lower threshold of 1 normalized transcript expression value (nTPM), defining that any receptor expressed below this threshold is not represented in the cell type/tissue. Of the 1,107 receptors, 26 were not found in the Human Protein Atlas and were therefore not included for further analysis. This MATLAB script has been deposited at https://github.com/Svensso n-Lab/danneskiold-samsoe2023.

### Construction of a ligand library

To construct a library of secreted proteins we collected all human entries listed as Secreted [SL-0243] under subcellular location [CC] in UniProt by 01-15-2023, n=3,845. From this pool, we kept reviewed entries (n=2,097), and entries longer than 16 amino acids (n=2,093). To limit computation, we also excluded sequences longer than 2,000 amino acids retaining 2,039 entries. We kept only entries with an annotated gene name, n=2,023. We further excluded immunoglobulins by excluding any gene with the name containing either “IGH”, “IGKC”, “IGKV”, “IGLC” or “IGLV” retaining 1,864 entries. In the case of duplicated gene names, we retained only the entry with the longest amino acid sequence retaining 1,862 secreted proteins in the final library (**Table S2)**.

### Predicting structures

We predicted ligand-receptor structures for each ligand against all receptors in the final libraries. We used ParallelFold^52^ in combination with AlphaFold 2.2.4 excluding the relaxation step, without template and using the reduced database to generate multiple sequence alignments (MSAs) for both ligands and receptors. To predict structures, we used Alphafold 2.2.4 either as stand-alone using precomputed MSAs and the same settings as above, or with ParallelFold predicting five models per ligand-receptor pair on the Danish National Supercomputer Computerome or Sherlock at Stanford University. Results were visualized with ChimeraX version 1.5^53^ using the best-aligning pair of chains between reference and match structure for comparing PDB entries with predictions. A list of PDB IDs for all proteins in the test set and UniProt IDs for all tested ligands is provided in **Table S1 and S3**. PDB files for predictions with a penalized ipTM value > 0.4 is available at https://purl.stanford.edu/bg124rf2339.

### Score prediction

The ipTM scores were extracted from the AlphaFold pickle files. Penalized ipTM was calculated by taking the median of available predictions and subtracting the median absolute deviation (MAD) as previously described^37^. The pDockQ score was calculated as previously described^18^.

### Generation of contact maps

Contact maps were generated using the bio3d package^54^ in R version 4.2.1 with secondary structures predicted using Stride^55^ using either structure from the PDB database or predictions of ligand-receptor pairs as inputs. Distances below 8 Å were considered contacts.

### Ligand and receptor characteristics

To determine the amino acid length of ligands that bind to single-pass or multi-pass receptors, we extracted accession numbers for all peptide receptor-ligand pairs in the two databases CellPhoneDB^29^ (111 pairs) and GPCRs^36^ (86 pairs). We used UniProt^56^ to determine whether receptors were single-pass or multi-pass membrane proteins and whether they were annotated as secreted as determined by subcellular location. We excluded any pairs where both or none were secreted (202 excluded), and any pairs without receptor classification (12 excluded). In addition, we extracted the ligand’s gene length and its signal peptide length. We only included ligands once, independent of the number of interacting receptors (224 excluded). We also excluded ligand-receptor pairs that did not have exactly one chain each (150 excluded). The final list includes the ligand amino acid lengths (gene length minus signal peptide length) of 130 multi-pass membrane proteins and 67 single-pass membrane proteins (**Table S5**). Of these, 86 are from the GPCRs database and 111 from the CellPhoneDB. The MATLAB script used to obtain and filter data is deposited at https://github.com/Svensson-Lab/danneskiold-samsoe2023.

### IgSF ligand-receptor predictions

We restricted the predictions to proteins that were included in ligand-receptor pairs that had 1) been reported both in Wojtowicz *et. al* and elsewhere, and 2) where structures for the ligand-receptor pairs reported in Wojtowicz *et. al* had not been released at the AlphaFold training date cut-off. The number of ligand-receptor pairs in this dataset was 83 (**Table S6**).

### Selection of orphan ligands

Selection of orphan ligand to test AlphaLigand was filtered as follows from 80 high-priority targets reported in Siepe *et al*^15^. Angiopoietin-related proteins were deemed unlikely to bind cell-surface receptors and removed (n=72). Ligands with a gene length < 100 were removed (n=68). Ligands not annotated as ‘secreted’ by Uniprot were filtered (n=59). Ligands without expression in the Expi293F expression system and controls were removed (n=50) (**Table S3**).

### Computational requirements

To reduce computational costs, we started by calculating MSAs for all receptors and ligands using up to 15 hours, 16 CPU cores and 8Gb RAM (**Figure S6A**). Since this step only has to be performed once, the calculation of MSAs significantly reduces computation time. Due to limited GPU access, we first ran predictions using only CPUs restricting it to a maximum of 10.5 hours, 12 CPUs and 64GB of memory (**Figure S6B**). In the, on average, ten percent of cases where structures did not complete prediction using CPU setting, we used GPUs with same settings as above except a maximum of 24h (**Figure S6C**). After running the eight test cases, we observed that no ligand-receptor pairs with a penalized ipTM value < 0.1 after the first prediction and < 0.2 for the second prediction obtained a final penalized ipTM > 0.5 (**Figure S6D-S6E**). For the prediction of receptors for orphan ligands, we therefore adapted AlphaFold to exit after the first predictions in cases where the ipTM value was below these values. As most of the predicted ligand-receptor structures have an ipTM value < 0.2 this also significantly reduces computational cost.

### Code availability

All codes to run the screen can be obtained at https://github.com/Svensson-Lab/AlphaLigand under the Apache License, Version 2.0.

### Contact for Reagents and Resource Sharing

Information and requests for resources should be directed to and will be fulfilled by the Lead Contacts, Niels Banhos Danneskiold-Samsøe and Katrin J. Svensson.

### Statistical analyses

Differences in ligand length for known ligand-receptor pairs were calculated using the Kolmogorov–Smirnov test in MATLAB. We used two-way ANOVA followed by Turkey’s test for multiple comparisons of differences in ipTM values between different input domains and IgSF binding status for ligand-receptor pairs in GraphPad Prism version 9.5.0 or R version 4.2.1, *p < 0.05, **p < 0.01, ***p < 0.001, ****p < 0.0001. A two-sided Wilcoxon rank sum test in R was used to compare differences in iPAE for IgSF pair binding status. All statistical analyzes were done on distinct samples without repeated measures. The Shapiro-Wilks test was used to test for normality.

---

**Figure S1.**
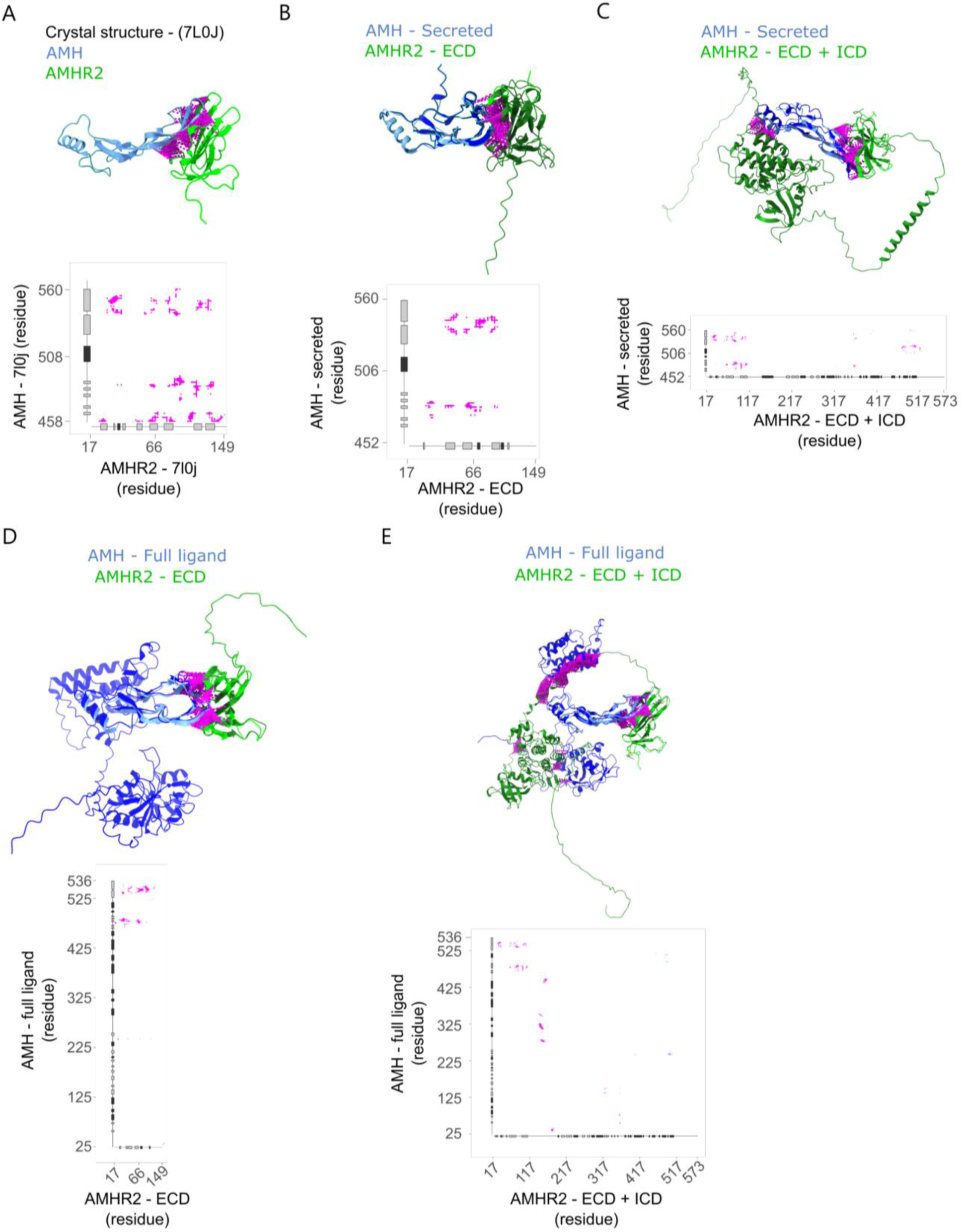
Binding prediction of single-pass receptor ligand complexes of AMH-AMHR2 using full or truncated sequences. (A-E) Structural binding prediction and corresponding contact maps for, light green: pdb database AMH, dark green: AF2 predicted AMH, light blue: PDB database AMHR2, dark blue: AF2 predicted AMHR2, purple: contacts (A) PDB entry, (B) secreted ligand and extracellular domain (ECD) of receptor, (**C)** secreted AMH and full receptor including intra (ICD), transmembrane (TCD) and ECD, (D) full ligand and ECD and (E) full ligand and full receptor. Tick labels indicate residue position in full length canonical protein (AMH accession= P03971, AMHR2 accession=Q16671).

**Figure S2.**
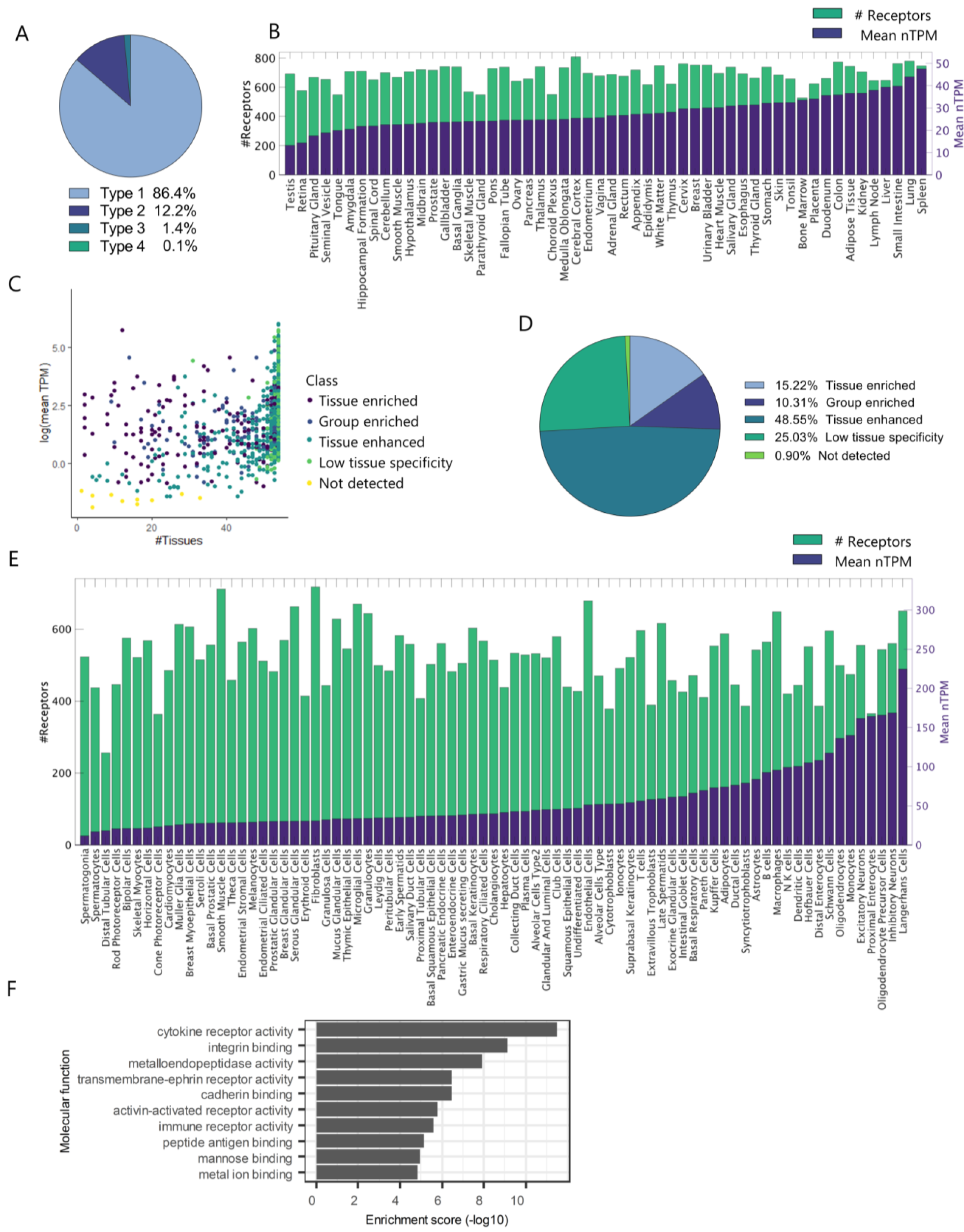
Composition of the single pass transmembrane receptor library. (A) Pie graph of tissue distribution of the receptors in the library according to the human protein atlas (HPA). (B) Receptor expression (mean nTPM) relative to the number of receptors (# Receptors) across tissues. (C) Receptor expression as mean normalized transcript per million (nTPM) relative to the number of tissues each receptor is expressed in. Each dot represents a receptor. Colors denote tissue distribution according to HPA. (D) Pie graph of the tissue expression of receptors in the library. (E) Receptor expression (mean nTPM) relative the number of receptors (# receptors) across cell types. (F) Classification of molecular functions for receptors in the library.

**Figure S3.**
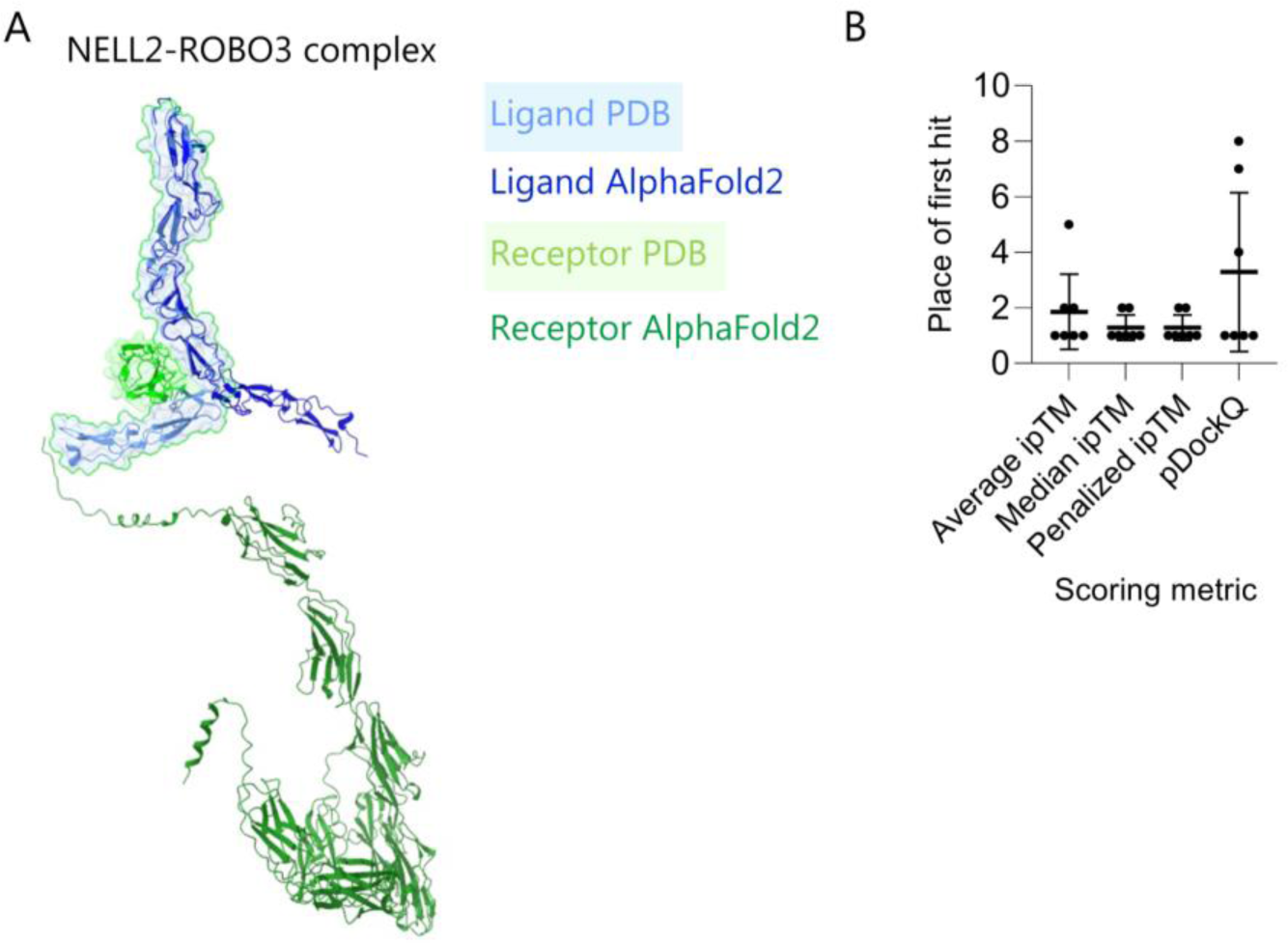
AF2 accurately predicts single-pass receptors for secreted ligands. (A) predicted structure of the NELL2-ROBO3 complex. Annotation as follows: light green cartoon and surface: PDB database receptor, dark green cartoon: AF2 predicted receptor, light blue cartoon and surface: PDB database ligand, dark blue cartoon: AF2 predicted ligand. (B) ranking of first hit when using the scoring metrics average ipTM, median ipTM, or penalized ipTM and pDockQ. Mean and 95% CI indicated.

**Figure S4.**
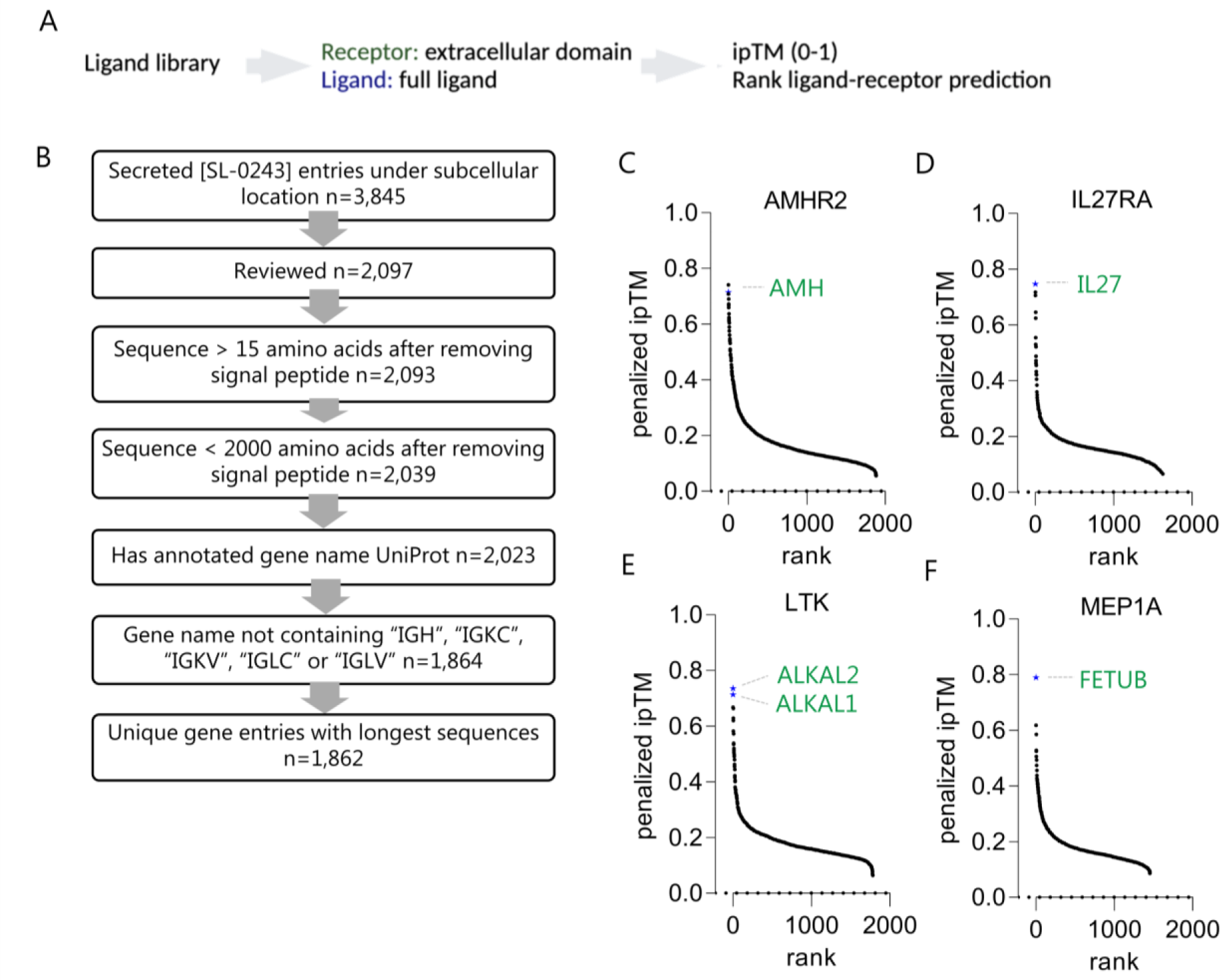
AF2 accurately predicts single-pass receptors for secreted ligands. (A) Approach and metric for scoring binding of single-pass transmembrane receptors against a ligand library. (B) Schematic of ligand library construction. 1: Extract human entries annotated as secreted [SL-0243] under subcellular location n=3,845. 2: Retain reviewed entries n=2,097. 3: Exclude secreted peptides/proteins with an extracellular domain shorter than 16 amino acids n=2,093 and 4: longer than 2,000 amino acids n=2,039. 5: Keep proteins with annotated gene names n=2,023. 6: Remove immunoglobulins with gene names either including “IGH, “IGKC”, “IGKV”, “IGLC” or “IGLV” n=1,864. 6: exclude duplicated gene names retaining entry with the longest sequence, n=1,862. Binding prediction of (C) AMHR2, (D) IL27RA, (E) LTK, and (F) MEP1A to the ligand library. Values are expressed as ranked penalized ipTM. The predictions are the median minus median absolute deviation of five independent predictions for each ligand-receptor.

**Figure S5.**
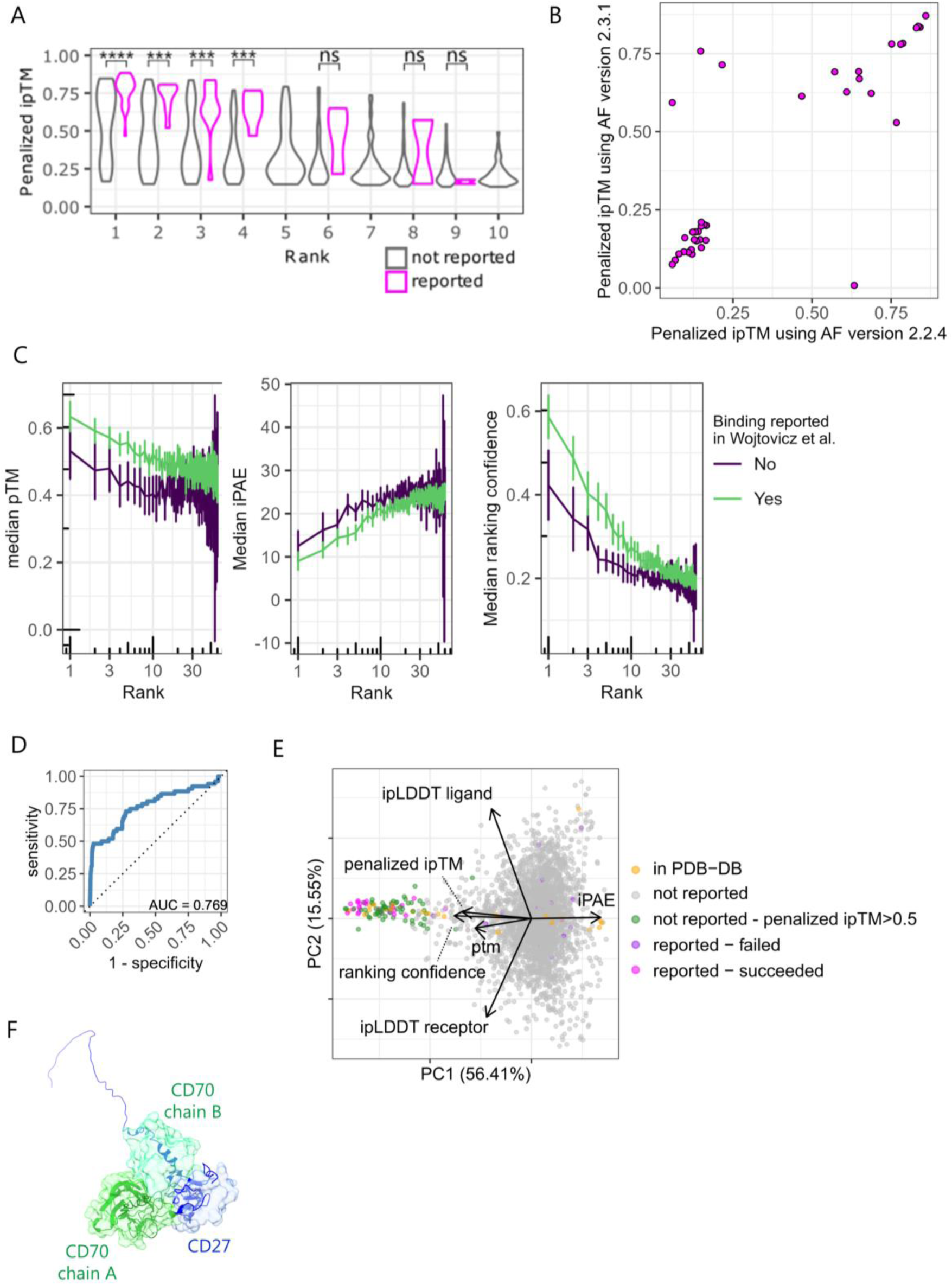
Ligands that fail prediction may be inferred from AF metrics. **(A)** Violin plot of relationship between rank and ipTM value for binding pairs in *Wojtowicz et al*. (n=2-79). One reported binding ranked fifth, seventh and tenth not shown. (B) correlation between penalized ipTM predicted using AlphaFold 2.2.4 and 2.3.1 of 40 ligand-receptor pairs that did not have released structures at the training date cutoff for AlphaFold version 2.3.0. (C) Median pTM, iPAE and ranking confidence as a function of rank for non-binding ligand-receptor pairs grouped by whether the ligand was successfully predicted (ipTM > 0.5) as binding to a receptor also determined to bind in Wojtowicz *et al*. (n=919-2430). Values are presented as mean ± 95% CI. (D) ROC curve using pairs identified in Wojtowicz *et al.* as ground truth (AUC=0.769). (E) principal component analysis (PCA) including indicated metrics from predictions in Fig. 4A. (F) best prediction of the CD27-CD70 heterodimer overlaid with partial crystal structure (7KX0) displaying compound interface, (green cartoon: predicted structure of CD70, blue cartoon predicted structure of CD27, green surface: CD70 crystal structure chain A, turquoise surface: CD70 crystal structure chain B, blue surface: crystal structure of CD27).

**Figure S6.**
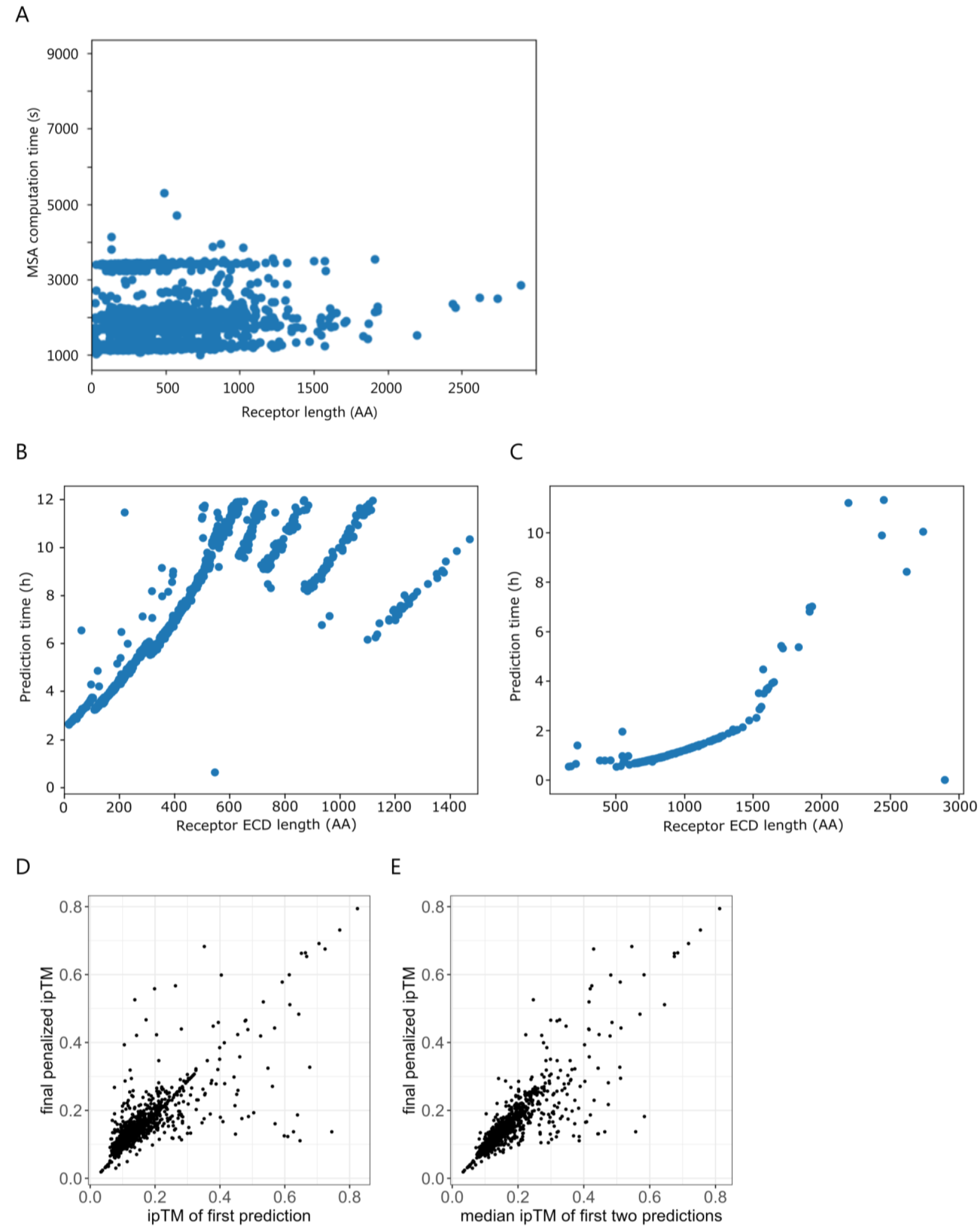
Computational cost and mitigation. (A) computation time needed for multiple sequence alignments (MSAs) by receptor length in amino acids (AA). (B) CPU time spent per receptor in library predicted with BMP10. (C) GPU time spent per receptor in library predicted with BMP10. (D) relationship between ipTM value in first prediction and final penalized ipTM value after five predictions for IL27. (E) relationship between median ipTM value in the first two predictions and final penalized ipTM value after five predictions for IL27.

## Supplementary Data Tables

**Table S1.** Information on the test set of ligand-receptor pairs with release dates after AlphaFold2 training date cut-off.

**Table S2.** Library of single-pass receptors included in receptor screen and secreted proteins in ligand screen.

**Table S3.** List of orphan ligands tested against the receptor screen. This panel was used for Fig. 5.

**Table S4.** Top-ranking receptor hits for 45 orphan ligands with at least one receptor with a penalized ipTM value > 0.5. List of orphan ligand-receptor pairs with missing predictions.

**Table S5.** List and data origin of ligand-receptor pairs either binding single-pass or multi-pass receptors used for Fig. 2d.

**Table S6.** List of genes in the IgSF superfamily used to test the performance of AlphaFold as screen related to Fig. 4.

## Notes

### Competing Interest Statement

The authors have declared no competing interest.

### Summary of Updates

Revised version has additional supporting data.

